# Head Direction Cells and Grid Cells as the Core Components of the Core-periphery Structure

**DOI:** 10.1101/2023.11.21.568067

**Authors:** Ruixin Qian, Yunxiang Chen, Tao Wang, Wei Wang, Feng Liu

## Abstract

Head direction cells and grid cells are neuron types defined by their regular firing patterns in standard experimental arenas. As technology advances, we can record extensive neuronal firing activity over time using electrophysiological or calcium imaging methods. The covariance matrix is a critical measure of this neural population’s discharge activity. We developed a method to identify a core-periphery structure in the covariance matrix, highlighting the central role of grid cells and head direction cells in firing correlations. This method effectively redefines these cell types in terms of firing activity correlations, with core nodes exhibiting a higher mutual information rate for spatial variables. Additionally, we developed a periodic spring network algorithm, which uses the covariance matrix alone to estimate the spatial phases of both head direction cells and grid cells due to their periodic properties. This approach offers a new perspective on utilizing the covariance matrix of the neural population to better understand and identify these specialized cell types, even when traditional firing pattern-based definitions are challenging to apply.

## Introduction

Head direction cells (HDCs) in anterodorsal thalamic nucleus (ADn) and grid cells (GCs) in medial entorhinal cortex (mEC) exhibit regular tuning curves tied to an animal’s head direction and location in its environment (Hafting et al., 2005; Taube et al., 1990). They can be easily identified based on the HD index and gridness score calculated from spatial tuning curves when the animal moves freely in a symmetric (circle or square) arena (Peyrache et al., 2015; Sargolini et al., 2006). In asymmetric environments, however, this classification method faces challenges as regular tuning curves can become distorted or disintegrate. Lattice-like firing fields of grid cells are deformed in trapezoid arena, sheared by walls, disintegrated in multi-compartment environments (Derdikman et al., 2009; Krupic et al., 2015; Stensola et al., 2015). On the other hand, the periodic grid cell firing fields are disrupted when the animal moves on a linear track (Yoon et al., 2016) and deviate from a regular hexagonal lattice on a circular track (Jacob et al., 2019; Low et al., 2021). Furthermore, HDCs and GCs get activated during sleep, but their spatial tuning curves are completely unavailable owing to a lack of environmental information (Gardner et al., 2019; Peyrache et al., 2015; Trettel et al., 2019). Given the ubiquity of asymmetric environments in daily life, exploring HDCs and GCs in these contexts holds particular significance.

Numerous methods have been introduced to address these distortions and disintegrations. Fitting an ellipse to the autocorrelogram of the rate map can detect slightly deformed and sheared grid patterns (Sargolini et al., 2006; Stensola et al., 2012), but it is ineffective in cases of severe disintegration. Grid cells were identified by assuming that their responses on linear tracks are equivalent to slices through a hexagonal lattice, with Fourier transformation used to analyze these responses (Yoon et al., 2016). The unique responses of GCs when varying the visual-motor gain on a linear track in virtual reality was suggested as a basis for classification (Low et al., 2021). But these assumptions call for more experiments to validate their universality. Overall, classifying neurons based only on tuning curves may not be a good choice, and accurately identifying HDCs and GCs under these experimental conditions remains a challenge.

Owing to the strict pre-defined criteria for neuron classification based on tuning curves, many cells remain unclassified, and unconventional yet crucial coding properties may be overlooked (Hardcastle et al., 2017). From the perspective of encoding and decoding, some neurons in the mEC exhibit mixed selectivity, i.e., their firing activities encode multiple spatial variables simultaneously, such as position, head direction and speed in a square arena (Hardcastle et al., 2017). This contributes to a high-dimensional representation of environments, contrasting sharply with the widely accepted scenario where grid cells establish a low-dimensional spatial code (Moser et al., 2014). Furthermore, decoding spatial variables using the entire recorded neural population consistently outperforms decoding using pools of strictly classified neurons alone (Stefanini et al, 2020), indicating that unclassified neurons also contain considerable spatial information. The inconsistency between decoding performance and conventional classification of neurons even in symmetric environments indicate systematic biases of the definition by spatial tuning curves. Notably, although the hexagonal firing fields of GCs and the head-direction tuning of HDCs are thought to serve functions as the universal metric and the compass of environmental space (Fyhn et al. 2007), the classifications only represent phenomenological descriptions of pre-defined spatial variables, which may explain the biases. Thus, it is essential to explain the biases, why neurons without regular spatial tuning could contribute to decoding, and explore a new classification paradigm, which reduces the biases.

Despite large variations in spatial tuning curves, the dynamic features of HDCs and GCs classified conventionally are maintained across various contexts. The joint activity vector of HDCs recorded simultaneously converges to invariant ring manifolds in phase space (Chaudhuri et al., 2019; Rybakken et al., 2019), while that of GCs revorded from the same module resides on a toroidal manifold (Gardner et al., 2022; Yoon et al., 2013). Notably, the manifolds are invariant across diverse environments, experimental blocks, and arousal states (Chaudhuri et al., 2019; Gardner et al., 2022). In accordance with the manifold representation, the correlations of firing activity between two HDCs (GCs) remain largely unchanged during sleep, and correlations of all cell pairs are preserved across states (Chaudhuri et al., 2019; Gardner et al., 2022; Gardner et al., 2019; Peyrache et al., 2015; Trettel et al., 2019; Yoganarasimha et al., 2006; Yoon et al., 2016). We are motivated to propose a classification approach relying solely on temporal firing sequences of neurons.

Here, we developed a novel method to distinguish HDCs in anterodorsal thalamic nucleus (ADn) and post-subiculum (PoS) and GCs in mEC from other neurons. A core-periphery (CP) network was constructed using activity correlations between simultaneously recorded neurons from the literature. Interestingly, conventionally classified HDCs (GCs) always constituted the core nodes, while the others, probably anatomically neighboring but insensitive to head directions (animal positions), formed the periphery nodes. On the basis of the CP architecture of unclassified neurons, HDCs (GCs) were also identified as the core neurons. The spatial information rate was used to assess the consistency of core neurons in spatial information, the distinctive characteristics of negative correlations between core nodes were established, and the spatial phases of HDCs (GCs) were determined using a spring network method. Grounded in the intrinsic dynamics of HDCs (GCs), our method surpasses conventional approaches relying on spatial tuning curves. Differences in classification results between our and conventional approaches offer insights into the dynamics and functions of HDCs and GCs.

## Results

### All recorded neurons indicating a core-periphery network

To illustrate distinct temporal features of HD and grid cells in comparison to other neuronal types, we analyzed four experimental datasets in the literature, all containing the activity of neurons and motion state of animals (Supplementary Table 1). The first dataset, referred to as *HD*, contains firing activities of neurons recorded from ADn and PoS of mice during both arousal (exploration in an open field) and sleep states (including rapid eye movement (REM) and slow wave sleep (SWS)). HDCs were identified using the HD index (Peyrache et al., 2015). The second dataset, named *GC-1*, comprises calcium signals from neurons in layer-II/III of mEC in mice during active movements within a square enclosure. Classic methods were employed to justify GCs, HDCs, border cells, object-vector cells, and others (Obenhaus et al., 2022). The third dataset, called *GC-2*, includes firing activities of neurons in mEC as mice navigated a linear track within a virtual reality (VR) environment. GCs and border cells were identified based on the observation that GCs were responsive to the discrepancy between visual and motor feedback in a 1D VR, whereas border cells remained stable (Low et al., 2021). Notably, the identification of GCs and border cells in this dataset differs from the classic method. The fourth dataset, denoted as *GC-3*, comprises firing activities of several hundred GCs in mEC when animals explored an open field arena or a wagon-wheel track, or during sleep, lacking firing activities of non-grid cells (Gardner et al., 2022). Collectively, HDCs and GCs have been classified according to their spatial tuning curves.

We first computed pairwise activity correlations among all cells recoded simultaneously in these datasets. As demonstrated in prior studies, the correlations between HDCs (GCs) displayed distinctive patterns; HDCs with similar preferred head directions or GCs with overlapping rate maps exhibited strong positive correlations, whereas those with opposite preferred directions or non-overlapping rate maps displayed negative correlations (Figure 1a and 1b). In contrast, correlations between HDCs and non-HDCs, GCs and non-GCs, and non-HDCs (non-GCs) did not demonstrate clear patterns. Notably, these patterns remained consistent across various contexts and behavioral states (Figure 1c and 1d). Thus, the correlation (at lag 0 s) distributions for HDC-HDC and GC-GC pairs presented long tails at both the negative and positive extremes, in contrast to the distributions for the other pairs (non-GC) (Figure 1e and 1f), as well as neuronal pairs belonging to different brain regions (Supplementary Figure 1). Together, HDCs (GCs) exhibit a consistent structure of activity correlation that sets them apart from other cell types.

**Figure 1.**
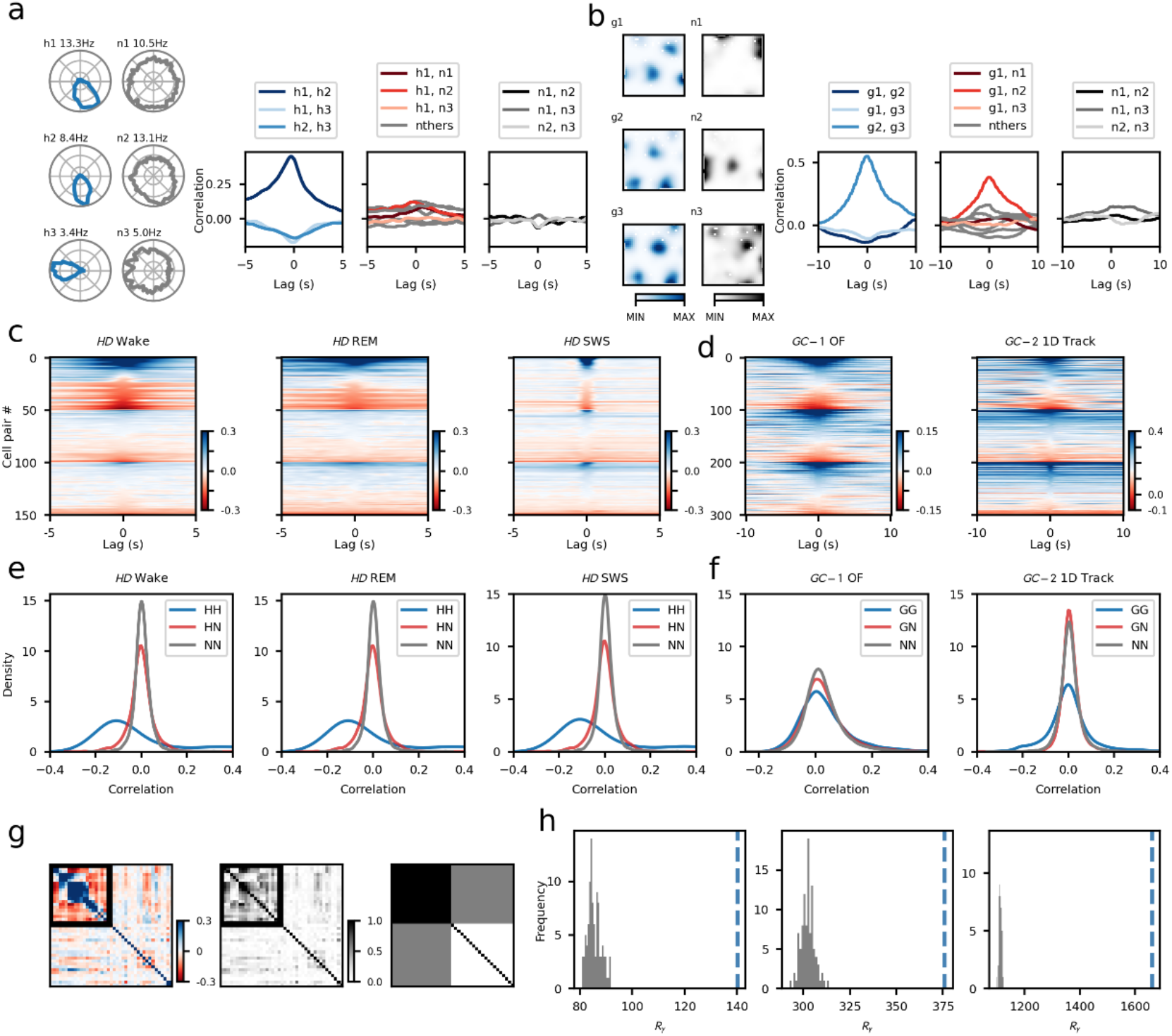
Activity correlation between different cells. **a.** Left: angular tuning curves of three HDCs (blue) and three non-HDCs (grey) from one session of wake state in *HD*. Right: corresponding activity correlations of HDC-HDC (left), HDC-non-HDC (middle, with only three examples color-coded), and non-HDC-non-HDC pairs (right). **b.** Left: spatial rate maps of three GCs (blue scale) and three non-GCs (grey scale) from one open-field (OF) session in *GC-1*. Right: corresponding activity correlations of GC-GC (left), GC-non-GC (middle, with only three examples color-coded), and non-GC-non-GC pairs (right). **c.** Color-coded correlation coefficient versus time lag for 50 examples of HDC-HDC (#0∼49), HDC-non-HDC (#50∼99) and non-HDC-non-HDC (#100∼149) from example sessions of wake state (Wake), rapid eye movement sleep (REM) and slow wave sleep (SWS) in *HD*. Each row shows a specific pair and is ranked within each pair type according to their correlation at lag of 0 s. **d.** Color-coded correlations for 100 examples of GC-GC (#0∼99), GC-non-GC (#100∼199) and non-GC-non-GC (#200∼299) from one example sessions of open field in *GC-1* and one example session of 1D track in *GC-2*. The same notation is used as in *c*. **e.** Kernel density estimation (KDE) plots for the correlations of HDC-HDC (*R*_HH_), HDC-other cell (*R*_HO_) and other-other cells (*R*_OO_) over all sessions of Wake, REM and SWS in *HD*. **f.** KDE plots for the correlations of GC-GC (*R*_GG_), GC-other (*R*_GO_) and other-other cell pairs (*R*_OO_) over all sessions of OF in *GC-1* (left) and over all sessions of 1D track in *GC-2* (right). **g.** Example of an activity correlation matrix of all recorded cells from a session in *HD* (left). The color from blue to red denotes the correlation strength from high to low. The positive and negative parts of the matrix are normalized separately, and then absolute values are taken (middle column). The color from black to white denotes the correlation strength from high to low. Schematic of the core-periphery structure (right), where black, gray and white indicate strong, weak and none connections. **h.** Distribution of shuffling results of 𝑅_!_, where the blue dashed lines indicate the experimental value.

These distinctive correlation patterns revealed a core-periphery (CP) network structure among all recorded neurons, with pairs of neurons connected by edges representing their activity correlation strengths. HDCs (GCs) served as the core nodes, while the remaining neurons contributed as the periphery nodes. The CP network is a fundamental mesoscale characteristic in network science, defined by densely connected core nodes and sparsely connected periphery nodes (Borgatti and Everett, 2000; Rombach et al., 2014). Within the mixed population of all neurons, HDCs (GCs) exhibited strong positive or negative correlations with each other, leading to dense connections, and thus constituted the core nodes; other neurons displayed weak correlations either with each other or with HDCs (GCs), acting as the periphery nodes (Figure 1g). Note that negative correlations were represented as absolute value because both large positive and negative correlations indicate strong interactions. To evaluate the adherence of the network to the CP structure, we calculated the core quality (*R*_γ_; Supplementary methods) using the matrix of all pairwise correlations. Across all datasets, a notably high *R*_γ_ was observed, surpassing the random results obtained through shuffling (Figure 1h). This conclusively demonstrate a distinct CP structure in the activity correlations among all simultaneously recorded neurons.

### Identifying HDCs and GCs by detecting the CP structure

The presence of a distinct CP structure in the activity correlation enabled us to propose a novel method for identifying HDCs (GCs) when spatial tuning curves are unavailable. This method involved assessing whether the covariance matrix formed by pairwise activity correlations of all recorded neurons adhere to the CP structure and identifying the core nodes as HDCs (GCs). To this end, we first calculated the activity correlation matrix for the mixed population (Figure 2a, Step I-II; Supplementary Methods). Next, we obtained the absolute values of the correlations (Figure 2a, Step III). Subsequently, we computed the core score for each node based on the CP structure, evaluating the extent to which the activity correlations align with CP networks of varying core sizes (Figure 2a, Step IV-V). A higher core score indicates denser connectivity among nodes.

**Fig. 2.**
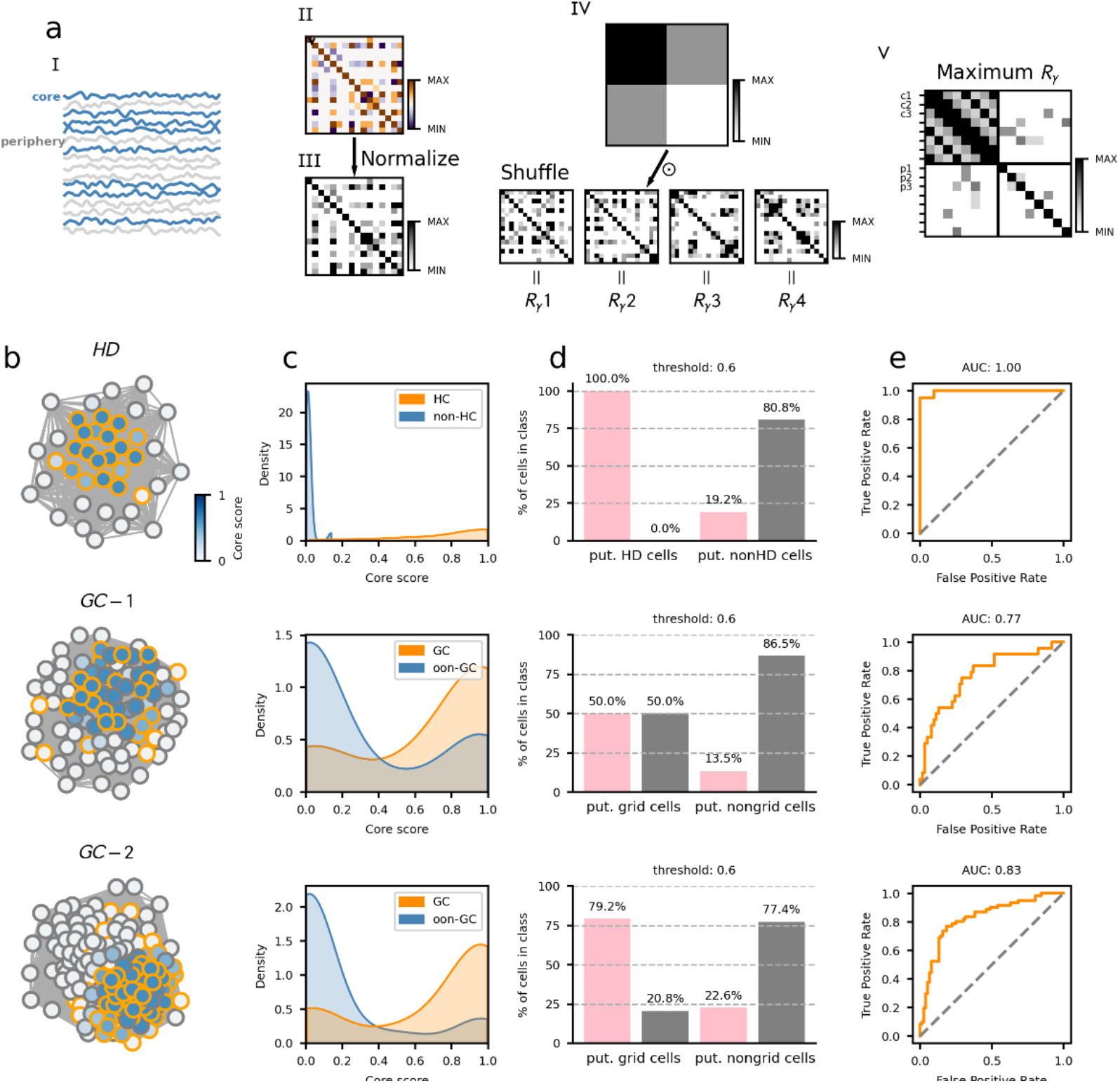
Detecting the core-Periphery structures. **a**. Flow charts for building the CP network (CPN). **I**, activities of recorded neurons; **II**, activity correlation matrix; **III**, normalized absolute values of the activity correlation matrix, where darker colors indicate larger correlation values; **IV**, shuffling the matrix to make it resemble a CP structure, performing element-wise multiplication and summation (𝑅_!_) between the shuffled matrix and the standard CP structure matrix (𝛽 = 0.5); **V**, acquiring the core score of each neuron. **b-e**. CPNs for three example sessions from three datasets. **b**. Core scores of all recorded neurons are labeled by the lightness of face color, with the lightest and darkest ones to be 0 and 1 respectively. Nodes with orange edges represent HDCs (GCs) with the classic method. The weight of lines connecting two nodes indicates their activity correlation strength. **c**. Probability density distribution of core scores for HDCs/GCs (orange) and other neurons (blue). **d**. Dichotomous results given a threshold of 0.6. In each putative category, HDCs and GCs classified classically are labeled in pink, while the others are labeled in gray. The bars display the fraction of cells in each putative category. **e**. Receiver operating characteristic (ROC) curve for the classification task, each giving an area under ROC curve (AUC). A higher AUC value indicates a stronger ability to distinguish HDCs (GCs) from others.

The results for three example populations from *HD*, *GC-1* and *GC-2* are illustrated in Figure 2b-e. A substantial number of neurons exhibited relatively high core scores, well consistent with the classic definition of HDCs (GCs) (Figure 2b); this suggests the effectiveness of the identification method, as neurons with higher core scores were more likely to be HDCs (GCs) as expected. To evaluate the method’s feasibility, we visualized the core score distributions of HDCs, GCs and other neurons. The distribution for HDCs (GCs) peaked around 1.0, while that for others peaked around 0 (Figure 2c). Based on a core score threshold of 0.6, recorded cells were categorized as putative HDCs (GCs) or others (Figure 2d). True positive and true negative results were prominently distinguished, demonstrating the accurate classification of HDCs (GCs) from other cell types (Figure 2d). Furthermore, the receiver operating characteristic (ROC) curve displayed an upper flank, and the area under the ROC curve (AUC) was remarkably large (Figure 2e). For the three examples presented, the AUCs were 1.00, 0.77, and 0.83, respectively, with a larger AUC indicating simpler classification. Additional session results are presented in Supplementary Figure 2. In conclusion, it is possible to distinguish HDCs (GCs) from other cell types even without spatial tuning curves.

### The CP method finding more putative grid cells than classic methods

Although the CP approach successively identifies HDCs and GCs, a performance disparity was observed in classifying GCs, with higher false negative and false positive rates compared with HDCs, leading to a lower AUC (Figure 3a). This means that HDCs exhibited dynamic independence and GCs did not. The primary reason could be the presence of diverse spatially tuned neurons within the mEC, including GCs, HDCs, border cells, and object-vector cells. During spatial navigation, these neurons engage in extensive interactions, potentially affecting the activity of GCs and blurring their dynamic independence. Conversely, the ADn and PoS predominantly contains HDCs, contributing to their higher degree of dynamic independence and fewer misclassifications. Overall, the increased interactions of mEC grid cells with other spatially-modulated neurons likely underlie the less distinct boundary between GCs and other cell types in terms of core score.

**Fig. 3.**
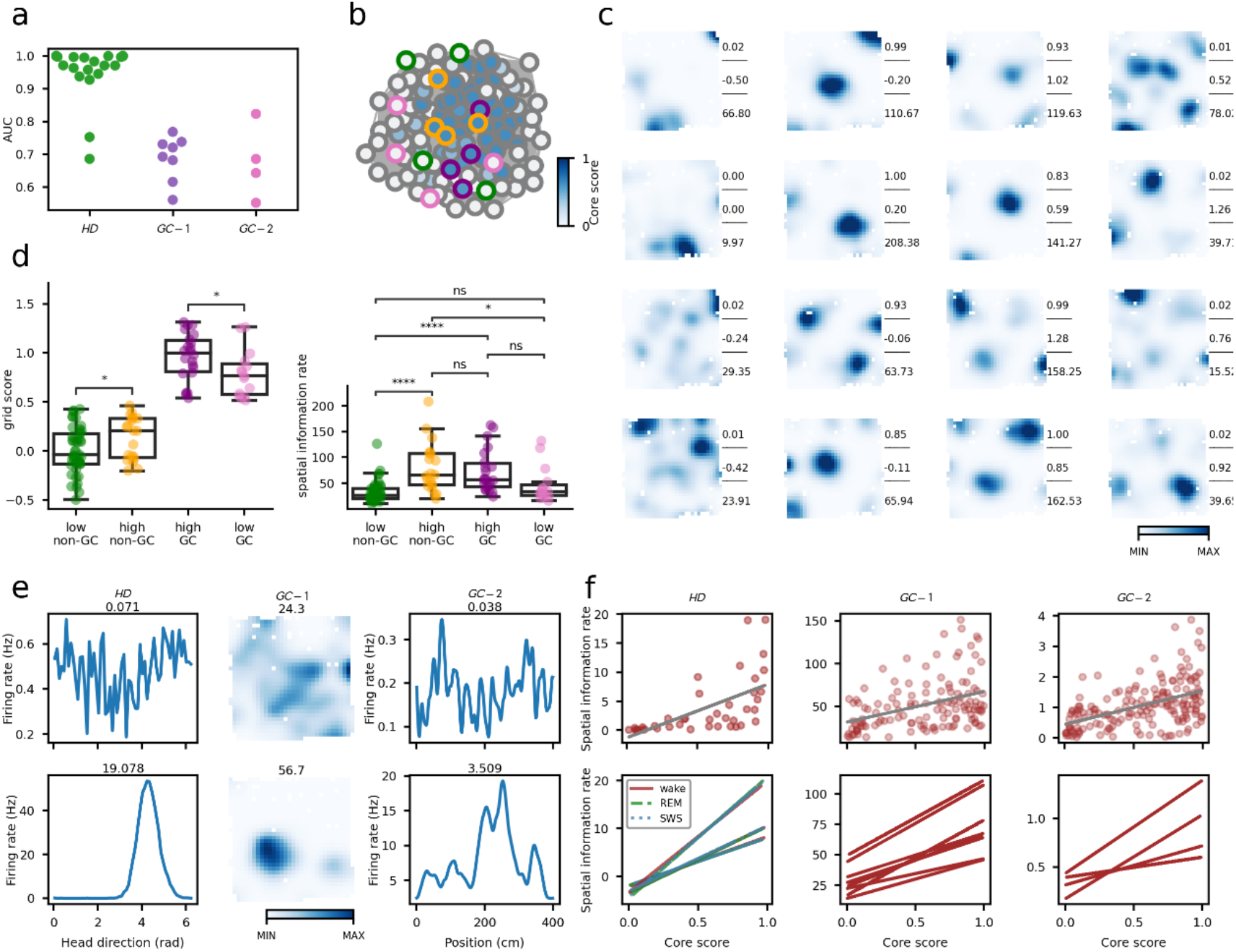
Statistical analysis of the classification results across three datasets. **a**. AUCs for all sessions across three datasets. **b**. Results for one session in *GC-1*. Core scores of all recorded neurons are labeled by the lightness of face color, with the lightest and darkest ones to be 0 and 1 respectively. Nodes with green, orange, purple, and pink edges are classified as non-GCs with low core score (< 0.2), non-GCs with high core score (> 0.8), GCs with high core score (> 0.8), and GCs with low core score (< 0.2). **c**. Each column shows rate maps of example neurons in the four categories in **b**. Values to the right are core score, grid score and mutual information (from up to bottom). **d**. Grid score (left) and mutual information (right) for the four different cell groups in **b** and **c**. *t*-test results are shown, with **** (p < 0.0001), *** (p < 0.001), ** (p < 0.01), * (p < 0.05), or ns (p > 0.05). **e**. The tuning curve and ratemap of six neurons with respect to head direction and position in three datasets, where the spatial information rate is low (above) and high (below). The spatial information rate is shown on each subplot. **f**. Correlation between the core score and spatial information rate of each neuron from three sessions in the three datasets, with **** (p < 0.0001). The bottom panel illustrates the correlation among 6 sessions in HD and all sessions in GC-1 and GC-2. For the six sessions in *HD*, distinct colors represent different states. For all sessions in *HD*, *GC-1* and *GC-2*, the p-value for Pearson’s correlation coefficient is less than 0.001.

The false negative rate may be attributed to the pronounced variability in spike timing of grid cells. For example, GCs often discharge much more vigorously than predicted by the mean firing-rate map or remain silent during specific traversals of firing fields (Hardcastle et al., 2015; Nagele et al., 2020). Thus, two GCs with similar spatial phases may occasionally exhibit a weak temporal correlation. Conversely, the occurrence of false positives suggests the existence of a neuronal type that exhibits strong dynamic correlation with GCs but is undetected by conventional methods. Intuitively, their firing-rate maps seem to resemble hexagonal lattices (Figure 3b and 3c).

To determine whether these false positive cells exhibit meaningful spatial tuning curves, we calculated the spatial information rate (*R*_SI_) between locations and neural activities (Supplementary methods). Cells with core scores above 0.8, including both GCs (true positive results) and non-GCs (false positive), displayed comparable *R*_SI_ levels, in contrast to cells with core scores below 0.2, regardless of whether they are GCs or not (Figure 3d). That is, non-GCs contain spatial information comparable to that of GCs as defined by classic methods. This implies that the traditional classification might be overly stringent and potentially susceptible to noise in neural dynamics or experimental recordings. Therefore, in contrast to the traditional definition of GCs based on spatial tuning curves, our method proves effective in identifying neurons which are highly likely to be GCs but are excluded by conventional approaches.

Moreover, we observed a positive correlation between the mutual information rate (*R*_MI_) and the core score (Figure 3e and 3f), which persisted across different sessions and states in *HD*. This positive correlation also held for the firing activity of mEC neurons on the one-dimensional track task (*GC-2*) across four sessions. These results suggest a direct link between the core score and a neuron’s encoding of spatial information, and this correlation extends across different states. In other words, the firing activity of neurons located at core nodes carries higher mutual information with the spatial variable every second. The core structure of the correlation matrix of firing activity for HDCs and GCs actually reflects their roles as the hub of information processing, indicative of functionally independent neuron types.

### Core score contributed by both symmetric positive and negative correlations

We further investigate the relationship between the core score and correlation symmetry by examining positive and negative connections separately. The connection symmetry evident in the figure can be represented by cliques, fully connected subsets in a graph (Figure 4a). Unlike edges, the formation of a clique necessitates not only strong positive or negative connections between pairs of neurons, but also the simultaneous establishment of these connections among groups of three, four, or even more neurons. Here we explored the relationships among symmetric negative correlation, grid score, and core score; the analysis parallels that for positive correlation. Consider three GCs with non-overlapping spatial phases (Figure 4d) and the firing activities of cell pairs all exhibited negative correlation. We modeled them as three nodes in a graph, with each edge representing the negative correlation (Figure 4d). We constructed a graph from a session in *GC-1*, retaining edges with negative connections below a 50% threshold. We then calculated two clique-related metrics: the number of cliques a node belonged to and the average node count in the cliques a node was involved in (see Supplementary methods). Clearly, the distributions of these two quantities were higher for GCs than for non-GCs, and higher for core nodes than for periphery nodes (Figure 4e). Owing to the even distribution of phases within the rhombic unit cell, GCs were more likely to form significant negative connections simultaneously among three or four GCs. Additionally, both symmetric positive and negative correlations contribute to the core score.

**Fig. 4.**
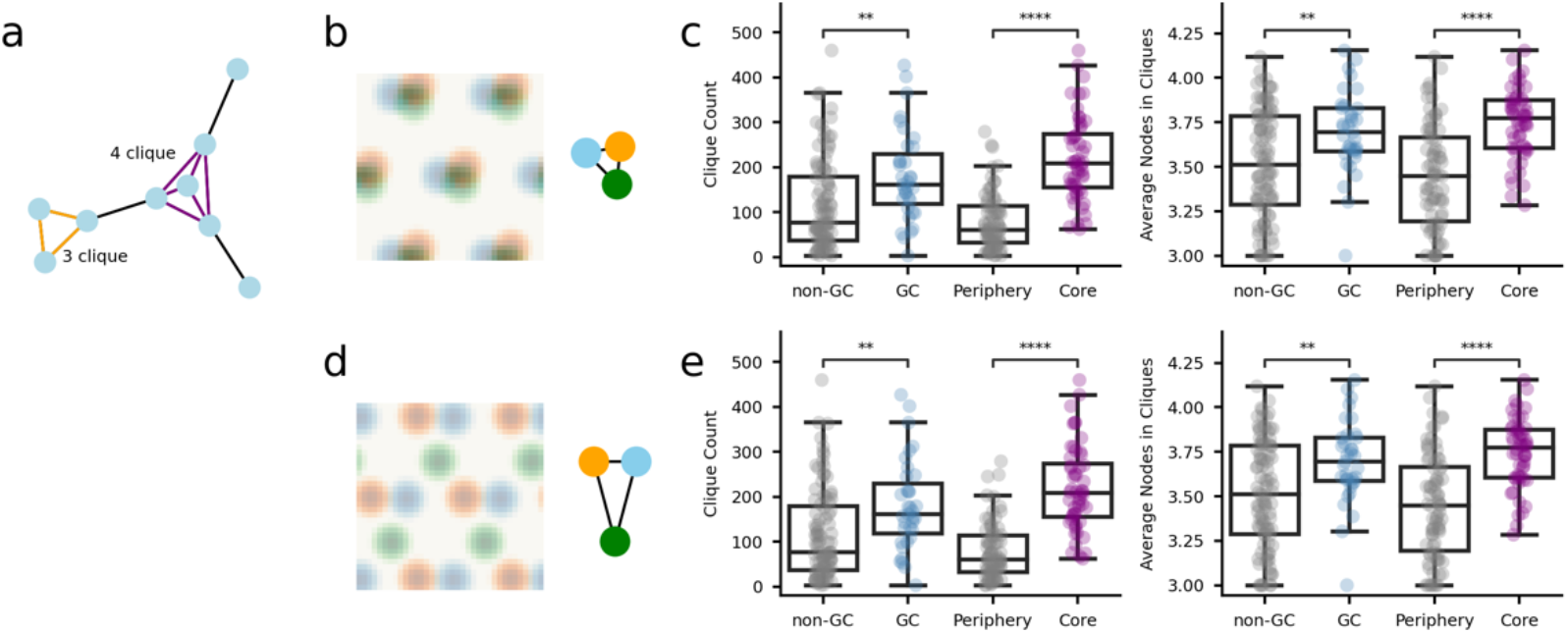
Both symmetric positive correlations and negative correlations contribute to the core score. **a**. Schematic of a network containing a 3-clique and 4-clique. **b** and **c**. Rate maps of three grid cells. The correlations of firing rates between these three neurons are positive/negative (left plot). In the graph with three nodes, each edge represents the positive/negative correlation between a pair of neurons (right plot). **d** and **e**. Neurons are classified into two groups based on their grid score: Grid cells (GCs) and non-grid cells (non-GCs), and categorized based on their core score: core nodes (core score > 0.6), and periphery nodes (core score < 0.6). The features related to cliques are calculated for the network, where nodes are connected by positive correlations (**c**) and negative correlations (**e**). The number of cliques each node participates in (where a clique consists of at least three nodes (left) and the average number of nodes in the cliques (right) for the four different cell groups. *t*-test results are shown.

### Relative spatial phases of HD and grid cells predicted by activity correlation

As mentioned earlier, the correlation structures of HDCs and GCs inherently reflect their spatial phase relationships (Figure 5a). Following the successful identification of HDCs and GCs within the mixed populations, we proceeded to explore whether it is possible to infer the preferred directions of HDCs and spatial phases of GCs from their activity correlation matrix. To improve our analysis, we introduced a simulated dataset termed *SD*, which presented activities of multiple neural populations along an experimentally recorded rat trajectory, including both HDCs and GCs (Kang et al., 2021). Similar to the experimental datasets described above, *SD* provided both spatial tuning curves and activity correlations for analysis. For the sake of simplicity and clarity, both the preferred direction of HDCs and the grid phase of GCs were collectively termed spatial phase. The spatial phases of HDCs fell within a range of 0∼2π, while those of GCs with the same grid scale were normalized in a rhombic unit with edges being 2π. This representation accurately mirrored the minimal repeatable structure of the grid lattice (Supplementary Figure 3).

**Fig. 5.**
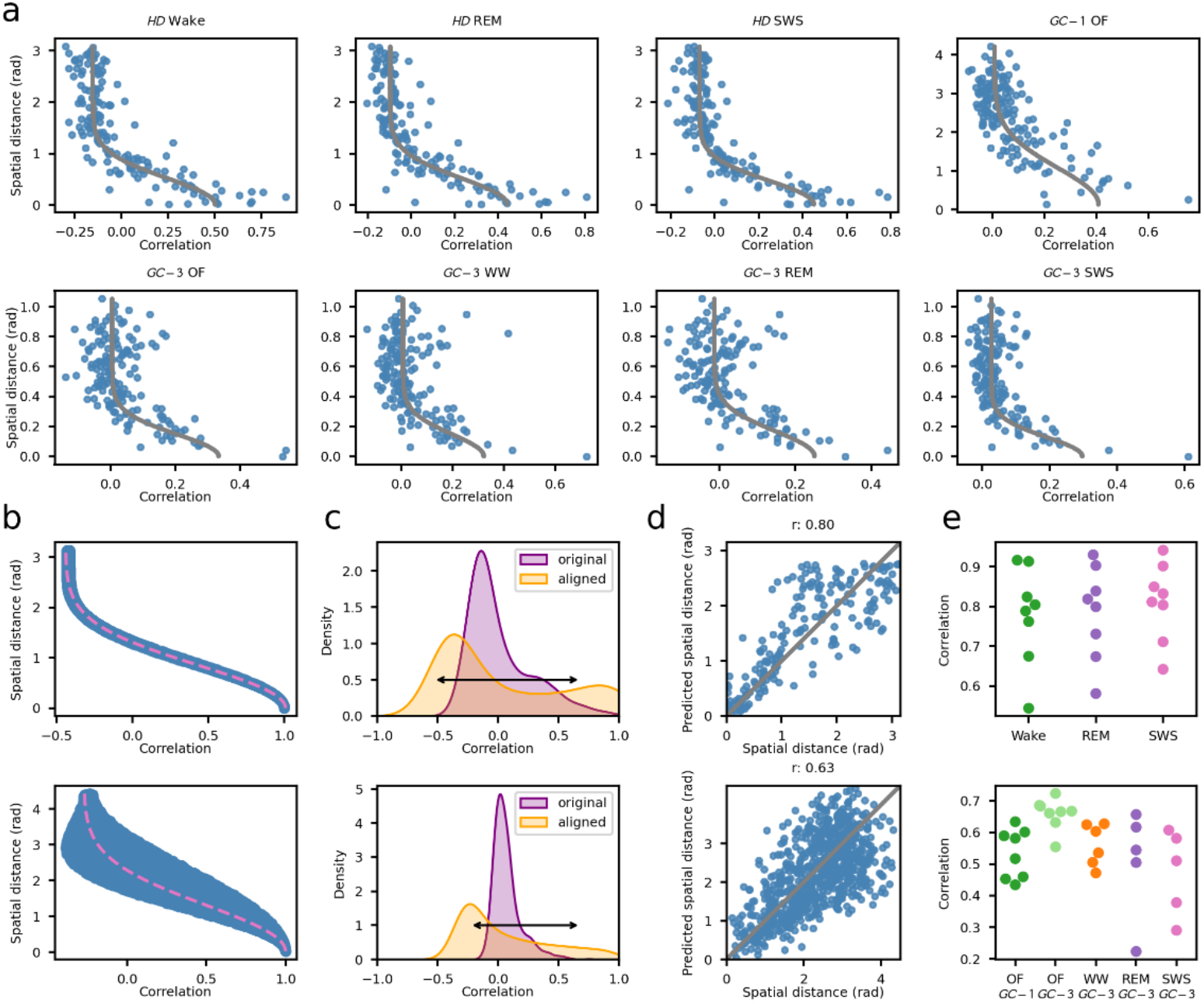
Predict the distance matrix based on the covariance. **a**. Relationship between the phase offset and activity correlation at lag 0 s for 150 HDC–HDC/GC-GC pairs (blue dots) from example sessions in different contexts in three datasets. The gray line indicates a fit to the data, which is the half of a Gaussian function. **b.** Numerical relationship between pairwise correlation and spatial distance. Blue dots denote the relationship for the *SD* dataset and the pink dashed line is a fit using the half of Gaussian function. **c.** The purple and orange curves illustrate the original and aligned pairwise correlation distribution for a session in the *HD* (top) or *GC-1* (bottom) dataset. The original experimental distribution is aligned to the simulated distribution before estimating the spatial distance using the function derived from the *SD* dataset. **d.** The predicted spatial distance is highly correlated with the true spatial distance for a session in the *HD* (top) and *GC-1* (bottom) dataset. **e.** Swarm plot of the statistical result illustrates the correlation between true spatial distance and predicted spatial distance across various sessions.

The activity correlation between two simultaneously recorded HDCs varied monotonically with their spatial phase difference in both *HD* and *SD*; an increase in phase difference notably led to a decrease in activity correlation (Figure 5a and 5b). Similarly, the activity correlation between two GCs in *GC-1*, *GC-3* and *SD* also depended on their spatial phase difference in the rhombic unit (Figure 6a and 6b). Although the angle of the phase difference of GCs impacted their activity correlation, the length of the phase difference played a dominant role (Supplementary Figure 4). This relationship was illustrated by a sigmoid function, albeit with a certain level of uncertainty in the phase difference axis (Figure 6b). Additionally, the distribution of activity correlation in *HD* was slightly narrower than that in *SD*, likely due to neural noise (Supplementary text and Supplementary Figure 5). Importantly, these relationships between HDCs (GCs) persisted across various recording sessions, experimental contexts, and datasets (Figure 6a). Conclusively, the connection between activity correlations and spatial phase differences is evident and straightforward for both HDCs and GCs.

**Fig. 6.**
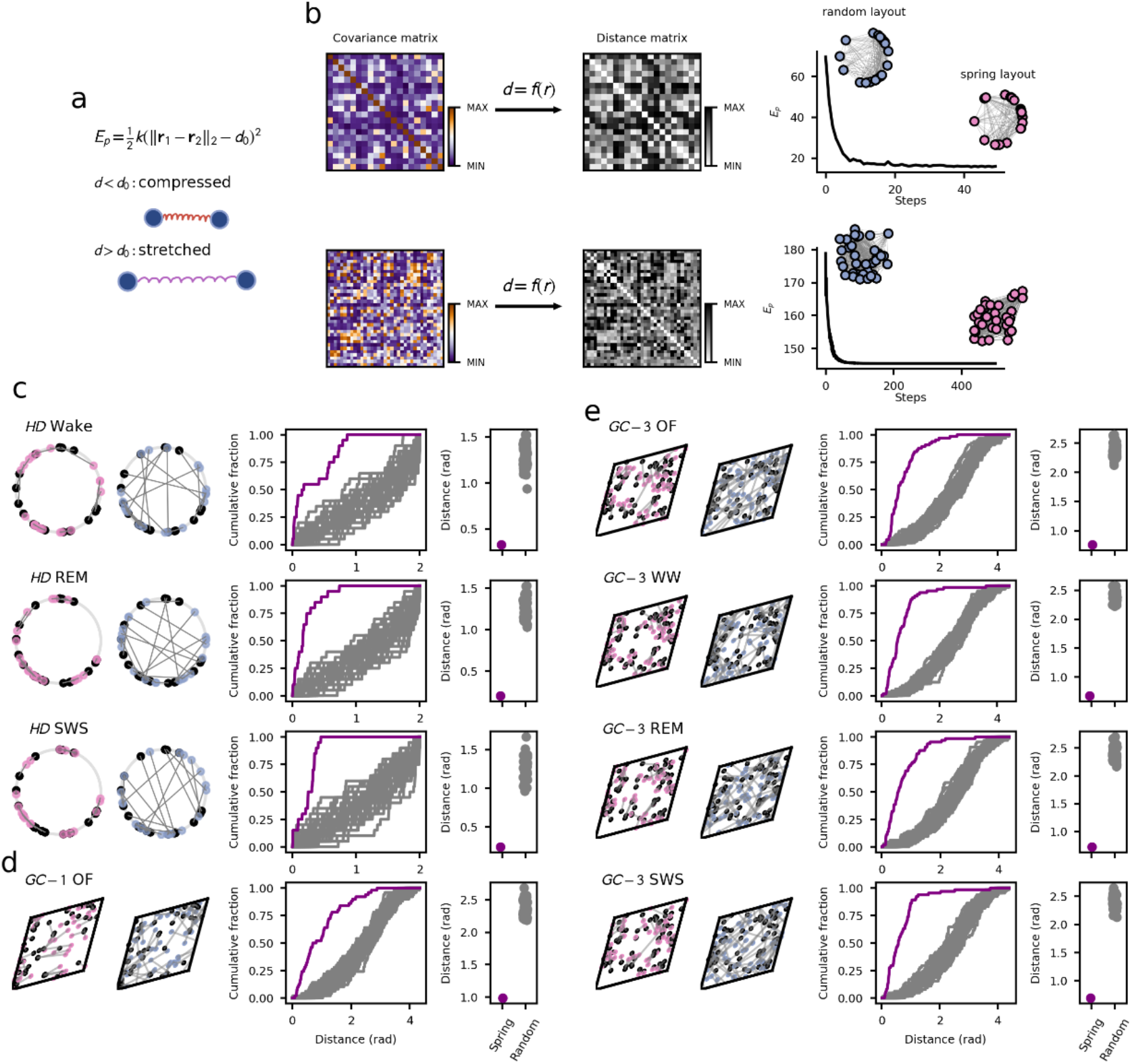
Attain preferred directions of head direction cells and spatial phases of grid cells. **a**. Two nodes are connected by a spring. **b**. The covariance matrix of a group of HD cells from a session of *HD* dataset and grid cells from a session of *GC-1* dataset (left column), contributing to the distance matrix (middle column). The initially randomly distributed neurons are iteratively arranged following the spring network method to minimize the potential energy (right column). **c-e**. Estimated spatial phases of three example sessions in *HD* (**c**), one example session in *GC-1* (**d**), and four example session in *GC-3* (**e**). **c**. Distributions of true spatial phases (black dots) and predicted spatial phases (pink dots) (the first column). Distributions of true spatial phases (black dots) and random spatial phases (blue dots) (the second column). Cumulative distributions of the distance between spatial phases (the third column), where the purple line corresponds to the spring layout and the blue lines correspond to 50 random layouts. Averaged distance between spatial phases for the spring layout and 50 random layouts (the fourth column).

Thus, it is possible to predict the spatial phase differences between two HDCs (GCs) through an inverse function relating activity correlation to spatial phase difference. Adopting the function derived from *SD* as a standard template, we first mapped experimental correlations to the simulated correlations. This mapping ensured alignment in the range between *SD* and experimental results and eliminating the influence of noise. Subsequently, we extracted the corresponding spatial phase differences (Figure 5c and 5d). Specifically, only the length of the spatial phase difference between two GCs was obtained. The predicted phase difference closely matched the actual one (Figure 5d), consistent across most sessions (Figure 5e). In sum, the phase differences between two HDCs (GCs) can indeed be predicted by their activity correlation.

Once the pairwise phase differences of HDCs (GCs) were determined, it became possible to establish the relative spatial phases for all cells. This involved arranging HDCs on a ring or GCs on a rhombus based on the obtained spatial phase differences. To achieve this, we utilized the spring network method (Supplementary method). Given the correlation matrix, the matrix of spatial phase differences (also called distance matrix), ***D*,** could be predicted as described earlier. The energy of the network was defined as 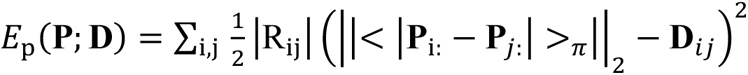, where *P*_i_ denotes the placement (phase) of node *i* and < 𝑎 >,≡ (𝑎 + 𝑚 𝑚𝑜𝑑 2𝑚) (Figure 6a). The goal was to minimize the system energy by iteratively adjusting ***P***, eventually positioning each node in a manner that was highly consistent with the distance matrix ***D***. The final placement, ***P***_F_, was determined by 𝑃_-_ = 𝑎𝑟𝑔𝑚𝑖𝑛_P_𝐸.(**P**; **D**) (Figure 6b). Notably, the graph was fully connected, meaning that the position of each vertex was influenced by all others. In this algorithm, |*R*_ij_| highlighted that the placement of a vertex was impacted significantly by the positions of its most positively and negatively related vertices.

This approach proved successful in accurately reconstructing the preferred directions of HDCs and spatial phases of GCs (Figure 6c-6e). Moreover, the spring layout remained stable across various conditions, validating this method and supporting the CAN dynamics of HDCs and GCs. Conclusively, the relative spatial phases of HDCs and GCs can be obtained from the activity correlation structures.

## Discussion

Traditional methods for identifying head direction cells and grid cells in rodents rely on their regular firing patterns in 2D environments. However, these methods struggle to pinpoint HDCs and GCs when rodents navigate in irregular environments or sleep, despite the persistence of neural firing activity correlations. Leveraging this characteristic and the core-periphery structure algorithm, we can identify putative HDCs and GCs and evaluate their functional independence using spatial information rate. Both positive and negative correlations contribute to neuronal classification. By applying the spring layout algorithm and analyzing the covariance of the neuronal population, we extract further information such as preferred directions of HDCs and spatial phases of GCs. Altogether, our results support CAN models of both HDCs and GCs. This method provides a valuable tool for studying neural coding in rodents.

The ever-increasing number of recorded neurons necessitates the development of algorithms for analyzing neuronal activity at a network and topological level. Here we present a network-based approach, where nodes represent individual neurons and edges signify pairwise correlation, a common modeling strategy in neuroscience (Belikovsky et al., 2015; Chang et al., 2019). A crucial underlying assumption is that the dynamic structure of HDCs and GCs remains largely invariant across different arousal states and contexts, as supported by numerous experiments (cite). Converting negative weights to their absolute values is essential for accurately identifying the core-periphery structure, where cores represent elements with high edge density. This adaption is necessary as both positive and negative correlations contribute to the core score. It is revealed that HDCs exhibit strong dynamic independence, in contrast to weak independence of GCs. Despite this, both HDCs and GCs demonstrate strong functional independence. Furthermore, the core-periphery structure in the mixed population aligns with the notion of functional independence of HDCs and GCs in earlier studies (Taube et al., 1990; Taube, 1995; Hardcastle et al., 2017; Obenhaus et al., 2022).

Our results provide strong support for the CAN model of both HDCs and GCs. The CAN necessitate denser connections among neurons within the CAN and sparser connections with neurons outside the CAN. Moreover, the preserved spatial phase relationship across arousal states and contexts confirms the invariance of CAN dynamics. From the perspective of the CP network, neurons within the CAN typically form core nodes, while those outside the CAN do not. Notably, the CAN model predicts that negative correlations between neurons are as important as positive correlations in determining core scores. This contrasts with neuronal assembles in other brain areas, which can also be analyzed using CP networks or similar methods (cite), but do not exhibit the similar level of negative correlation. In summary, our method provides a novel perspective on CAN dynamics, highlighting the important role of negative correlations between neurons.

In addition to our spring network method, circular coordinates (de Silva et al., 2011; Perea, 2018) can be used to predict the phases of individual neurons in state space by leveraging 1-cocycles extracted through persistent cohomology (Cohen-Steiner et al., 2010; de Silva et al., 2011; Edelsbrunner and Morozov, 2017). The phase in state space is highly correlated with the phase in the physical space (Rybakken et al., 2019) (Gardner et al., 2022). However, persistent cohomology is highly sensitive to noise and requires a large number of recorded head direction cells and grid cells, limiting its applicability to stringent conditions. The preprocessing steps required for persistent cohomology, such as dimensional reduction using PCA and subsampling points in the state space, are computationally intensive. By contrast, our method requires fewer neurons and less computation, while still achieving accurate predictions of the spatial phase of each neuron. This enhances its generalizability and practicality.

Our methodology could be improved in several manners. First, the current approach of calculating the core score of the covariance matrix assumes that neurons within a continuous attractor have stronger temporal connections. However, other crucial features, such as Fourier coefficients of neuronal firing patterns, could also be used to distinguish neurons. To address this limitation, we could develop a classifier that automatically clusters neurons and identifies defining features for various neuronal populations based on their firing activity. Second, to effectively distinguish HDCs and GCs from other neural types, we need record a considerable number of these cells in the mixed population. If no enough HDCs (GCs) are present, the network may not exhibit a clear core-periphery structure.

